# A Rapid COVID-19 RT-PCR Detection Assay for Low Resource Settings

**DOI:** 10.1101/2020.04.29.069591

**Authors:** Arunkumar Arumugam, Matthew L. Faron, Peter Yu, Cole Markham, Season Wong

## Abstract

Quantitative reverse transcription polymerase chain reaction (RT-qPCR) assay is the gold standard recommended to test for acute SARS-CoV-2 infection. It has been used by the Centers for Disease Control and Prevention (CDC) and several other companies in their Emergency Use Authorization (EUA) assays. RT-qPCR requires expensive equipment such as RNA isolation instruments and real-time PCR thermal cyclers, which are not available in many low resource settings and developing countries. As a pandemic, COVID-19 has quickly spread to the rest of the world. Many underdeveloped and developing counties do not have the means for fast and accurate COVID-19 detection to control this outbreak. Using COVID-19 positive clinical specimens, we demonstrated that RT-PCR assays can be performed in as little as 12 minutes using untreated samples, heat-inactivated samples, or extracted RNA templates. Rapid RT-PCR was achieved using thin-walled PCR tubes and a setup including sous vide immersion heaters/circulators. Our data suggest that rapid RT-PCR can be implemented for sensitive and specific molecular diagnosis of COVID-19 in situations where sophisticated laboratory instruments are not available.

## Introduction

Quantitative reverse transcription polymerase chain reaction (RT-qPCR) assay is the gold standard recommended to test for acute SARS-CoV-2 infection.^1–4^ It has been used by the Centers for Disease Control and Prevention (CDC) and several other companies in their Emergency Use Authorization (EUA) assays.^5^ Despite its established performance in sensitivity and specificity, RT-qPCR requires expensive equipment such as RNA isolation instruments and real-time PCR thermal cyclers, which are not available in many resource limiting settings.

As a pandemic, COVID-19 has quickly spread to the rest of the world. Many underdeveloped and developing counties do not have the means for fast and accurate COVID-19 detection to control this outbreak. Therefore, any development toward rapid, sensitive, and affordable RT-PCR assays could help advance the diagnosis and thus limit the spread of COVID-19.

Our approach is to update the “archaic” method of hand-transferring reaction tubes through a series of water baths. Two water baths (one for denaturation and one for the reverse transcription and then the annealing/extension steps) were made using food storage plastic containers heated by sous vide immersion heaters. These heaters are easy to use and have precise temperature control and circulating functionality. The PCR tubes were then shuttled between water baths using a servo motor operated arm controlled by a Raspberry Pi-based device.^6–8^

Our device costed ~$300 to build, whereas other PCR machines typically cost thousands of dollars. In addition to its low price tag, the unit can process up to 96 samples in one run. It is also extremely fast, as 40 cycles of TaqMan probe-based PCR can be completed in 25 minutes using conventional polypropylene PCR tubes. Using thin-film PCR reaction tubes (e.g., Cepheid SmartCycler or glass capillary tubes), a 40-cycle RT-PCR reaction can be completed in as little as 12 minutes (2 minutes RT and 10 minutes PCR with 15 seconds per cycle). Compared to most benchtop thermal cyclers, this is a significant improvement in speed. While there was no real-time fluorescence detection during the run, the results of the probe-based RT-PCR assays were examined using the end-point PCR approach. The fluorescence signal of the tubes after amplification was examined by placing the tubes on top of a blue LED gel-viewing box and viewed through an amber or orange colored filter. A cell phone camera was used to record the signal difference between the PCR tubes.

We tested this rapid water bath RT-PCR approach using untreated or heat-inactivated samples directly added to one-step RT-PCR master mixes without an RNA extraction step. We have also developed a 3-minute rapid extraction step using a magnetic particle-based method to produce RNA templates. The results are described in the next section.

## Results/Discussion

### Ability in maintaining temperatures at denaturation and annealing/extension temperatures for RT-PCR

The sous vide immersion heaters we used are capable of heating water up to 99°C. This means they can be used to make an environment suitable for denaturing nucleic acid and amplicons. We used two 6-quart clear food storage containers (5.7 L) as water baths and we mounted each immersion heater to the side. The immersion heater can warm the water to 60°C in less than 10 minutes from room temperature water. A temperature data logger probe was immersed in the water to create a temperature plot (**Figure 1**). Following the initial heating, the immersion heater was left in the container for another 30 minutes to observe for its ability to maintain water temperature. After 30 minutes, the data logger showed that the water bath temperature remained at a steady 60°C, with only 0.1°C of variation.

**Figure 1.**
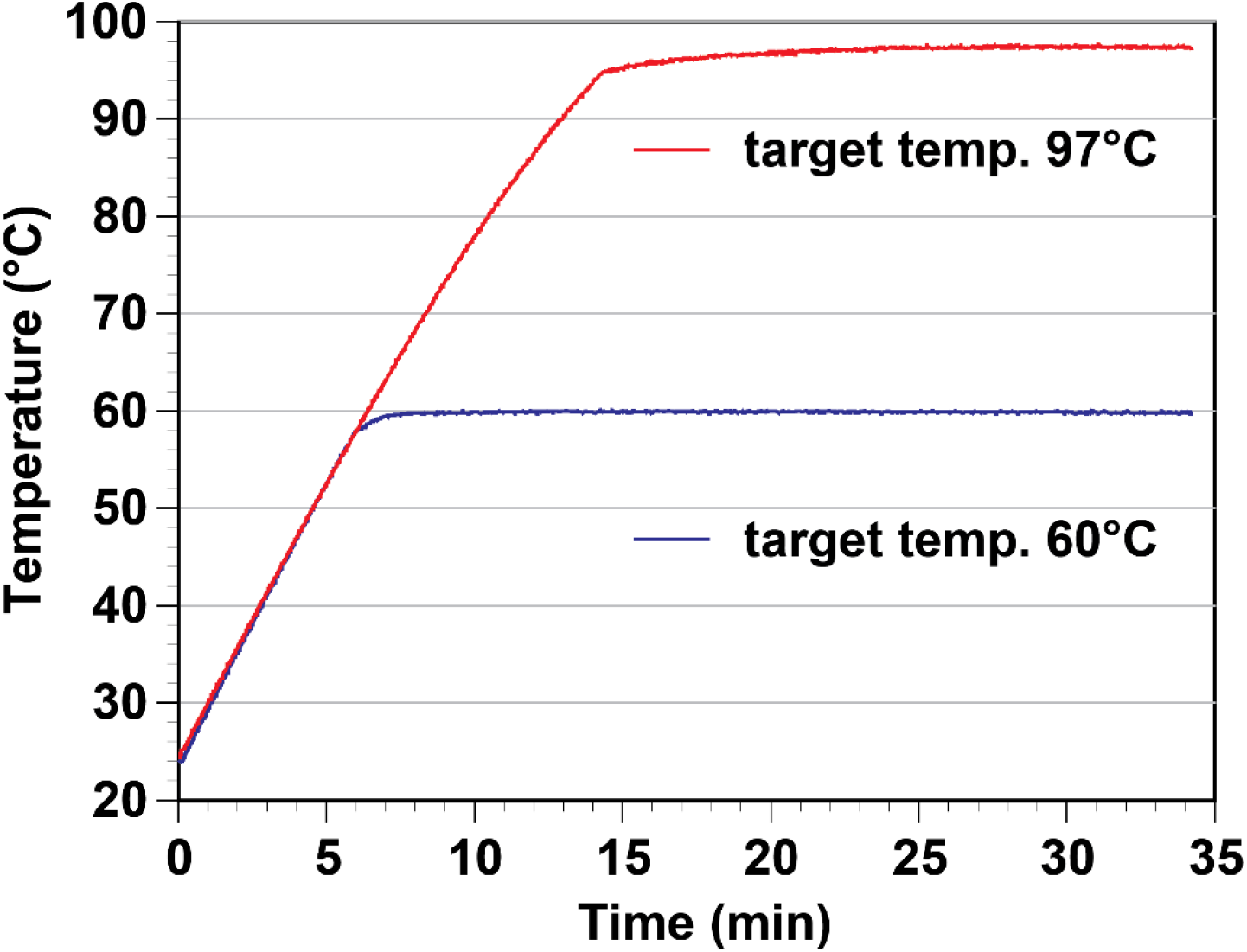
A sous vide immersion heater can maintain water bath temperatures at denaturation and annealing/extension temperatures steady enough for thermal cycling reactions.

In another test, room temperature water was heated to 97°C from a room temperature of 25.4°C. After 20 minutes, the immersion heater successfully heated the water to a steady 97°C. As expected, the heater successfully maintained a temperature of 97°C with only 2°C of variation. These two results confirm that the sous vide immersion heater can maintain steady heat long enough for the denaturation and annealing/extension steps of RT-PCR. The only limitation we expect for this set up is when using it in high altitude locations where boiling temperature is lower than 95°C. Lower denaturation temperature setting would be needed, along with longer denaturation time.

### RT-qPCR analysis of ten COVID-19 positive clinical samples

To determine the relative viral load of the clinical samples, 60 μL of the specimen was extracted using an automated system (Promega Maxwell) and run on a commercial real-time thermal cycler. Three μL of the templates were used in each RT-qPCR reaction. The qPCR threshold cycle (Ct) values of these samples are listed in **Table 1**. It took the commercial cycler 1 hour and 22 minutes to complete the reaction. Furthermore, to test the need for RNA extraction, 3 μLs of each media was spiked in 17 μL of the master mix. The results show that untreated media samples can produce qualitative PCR results matching those performed using extracted RNA templates.

**Table 1.**
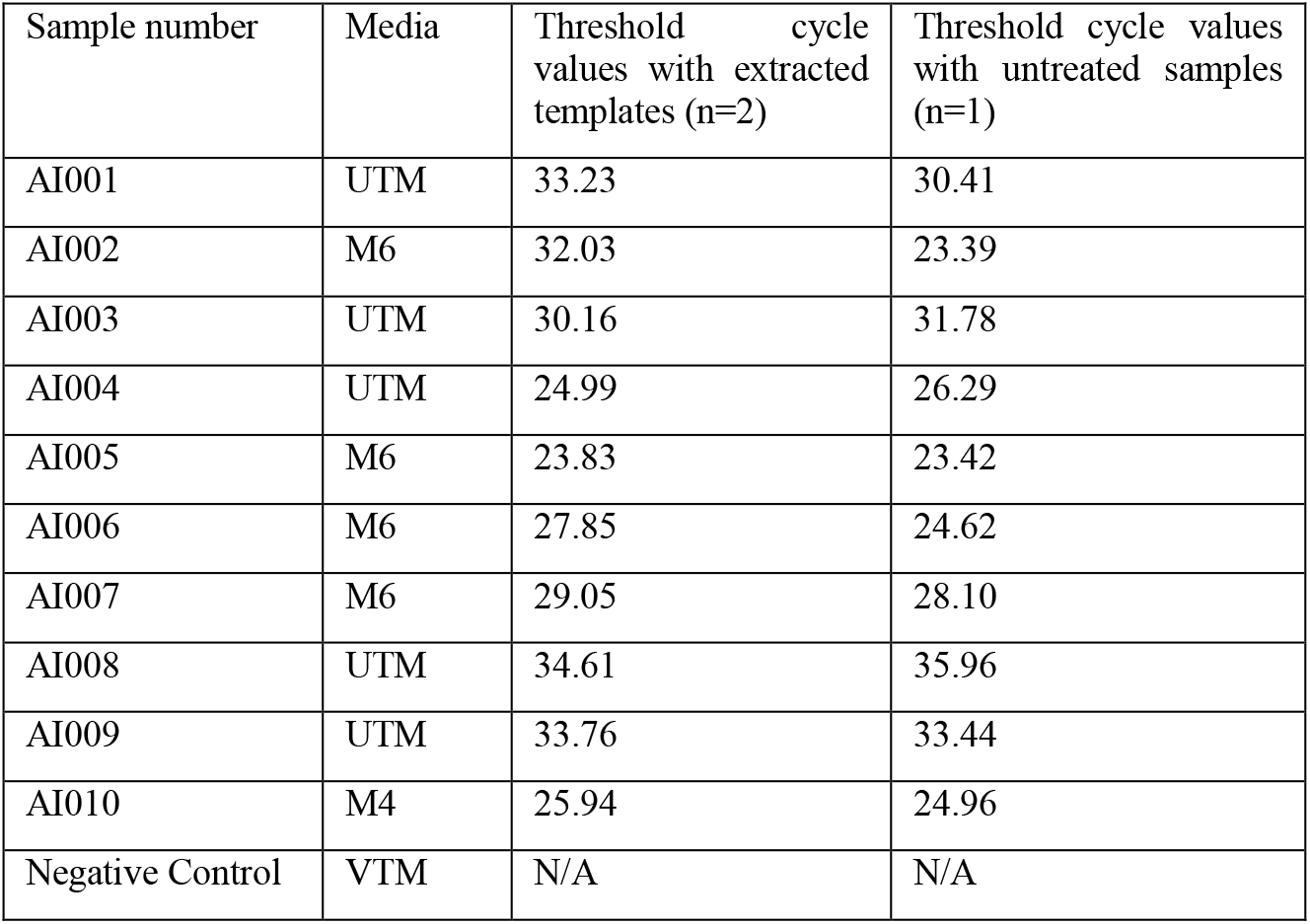
Threshold cycle values of our COVID-19 positive samples

### Rapid RT-PCR detection of SARS-CoV-2 with N1, N2, and RNase P reactions using water baths

To test for SAR-CoV-2 using water baths, we prepared three singleplex PCR reactions, each targeting N1, N2, and RNase P. Extracted templates from COVID-19 positive clinical specimens and contrived negative samples were tested using water bath-based RT-PCR. The RT-PCR run was completed in 12 minutes. The picture of the PCR tubes after 40 cycles shows that we can use a blue LED gel box and cell phone camera to determine the test results (**Figure 2**). In a COVID-19 positive sample, all three sample tubes had increased fluorescence signal after the reaction. In a COVID-19 negative sample, only the RNase P tube had increased in fluorescence intensity.

**Figure 2.**
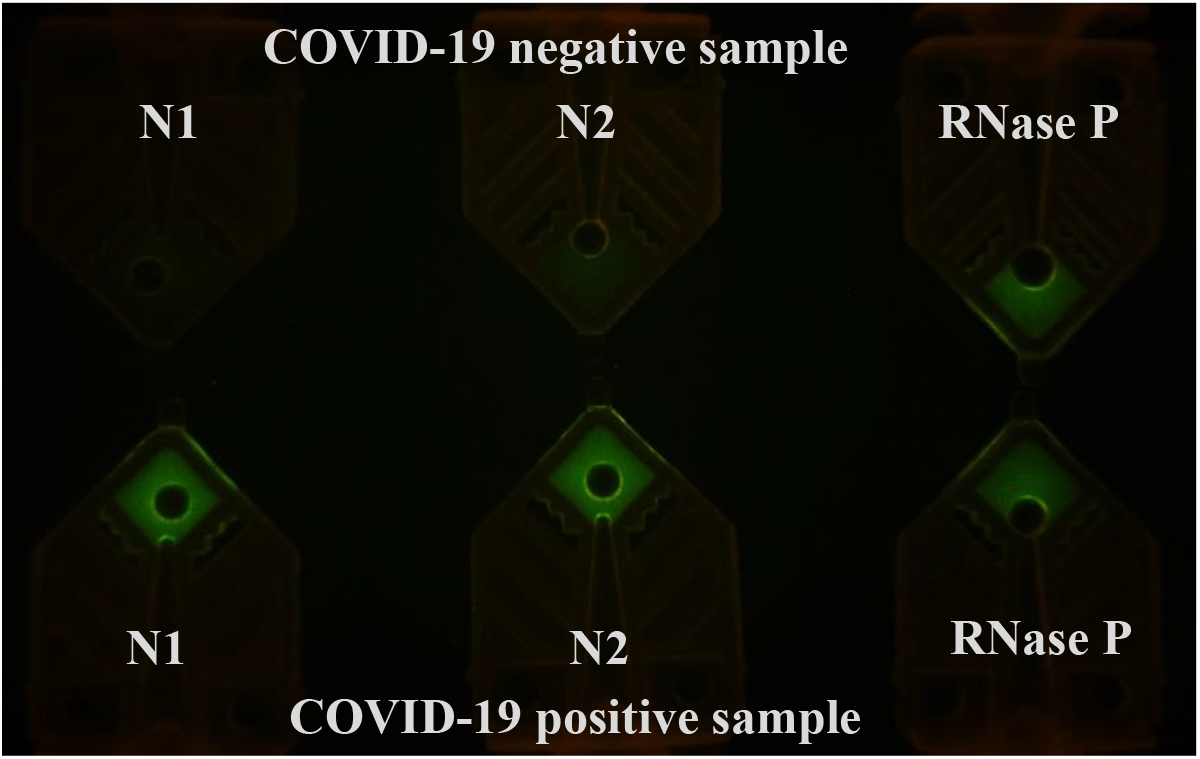
N1, N2, and RNase P in thin-walled PCR tubes after a RT-PCR run using 3 μL of extracted templates. Total run time was 12 minutes (90s/30s/40x(6xs/9s)).

We repeated the work using regular PCR tubes. While the reaction took longer to complete (26 minutes total), the results matched those of the thin-walled PCR tubes (**Figure 3**).

**Figure 3.**
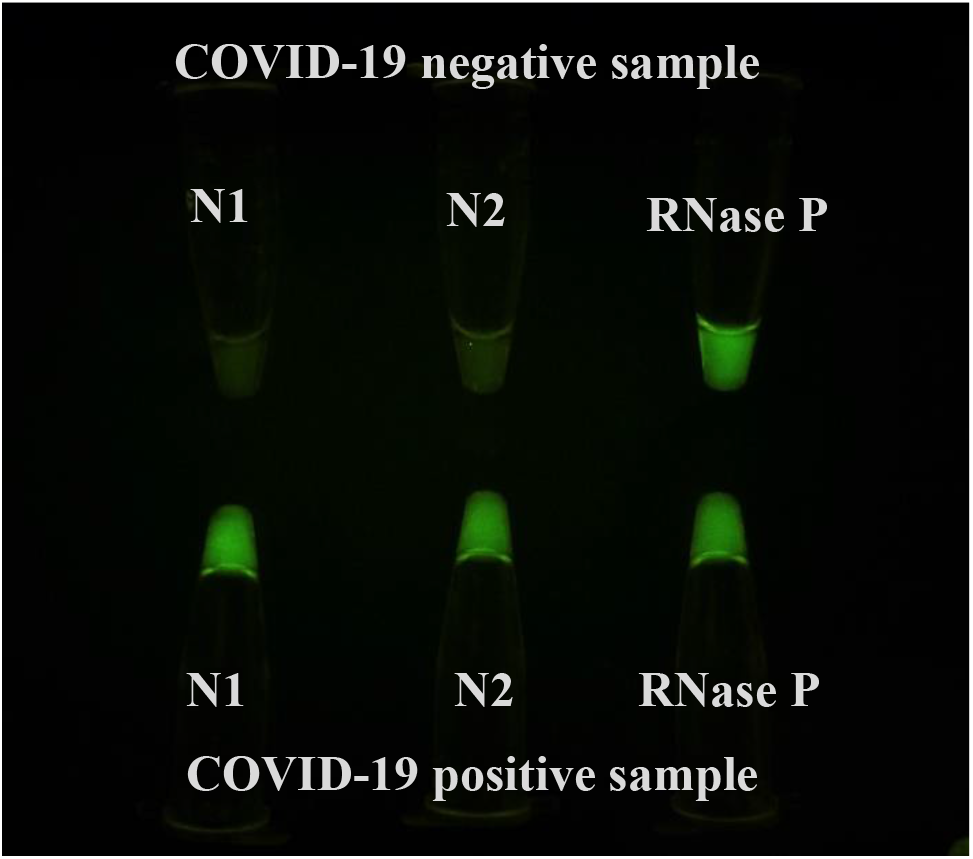
N1. N2, RNase P regular PCR tubes after RT-PCR reaction using 3 μL of extracted templates. Due to slower heat transfer, longer incubation time was needed. Total run time was 30 minutes (120s/30s/40x(15s/25s)).

### Rapid RT-PCR detection of SARS-CoV-2 in untreated clinical samples

The sample preparation step is generally time-consuming, regardless of whether it is done manually or automated. In addition, there was a shortage of the recommended viral RNA extraction kits needed for the Centers for Disease Control and Prevention (CDC) RT-qPCR assay to diagnose SARS-CoV-2. We investigated the feasibility of omitting the full RNA extraction step to expedite the test without significantly impacting the test’s sensitivity. Our study corroborates the findings from others who have successfully performed COVID-19 RT-qPCR reactions in a benchtop real-time thermal cycler by simply adding a few microliters of the unprocessed sample in transport medium directly into the RT-qPCR assay master mix.^9–12^ The data presented below suggests that it is possible to skip the RNA extraction step and use rapid RT-PCR to deliver fast COVID-19 testing using minimal equipment. While this approach may not be desirable for use in developed countries, it will find applications in many remote regions inside underdeveloped and developing countries where extraction devices and PCR thermal cyclers are not commonly available.

We first tested whether we could use untreated samples directly by spiking them into RT-PCR master mix to expedite the RT-PCR test (targeting N1) by skipping the extraction step. We performed the initial reactions with a protocol of 90 seconds RT, 30 seconds of reverse transcriptase inactivation and hot start for the PCR. This was followed by 40 cycles of 6 seconds of denaturation and 9 seconds of annealing/extension steps. Afterwards, photos were taken of the tubes illuminated on a blue LED gel box with an amber viewing filter (**Figure 4**). By comparing the tube intensity with the negative control, we were able to call 7 out of 10 of the samples as N1 positive (samples 2, 3, 4, 5, 6, 7, and 10). Sample 3 was a weak positive. The signal intensity for samples 1, 8, and 9 was not significantly different from the negative controls shown in the photo. These four untreated samples (1, 3, 8, and 9) all produced Ct over 30 when tested by RT-PCR (Table 1). We also speculate that the sensitivity of this run using untreated samples was low (70%) because we only spent 90 seconds on reverse transcription. The inhibitors in the samples can affect the cDNA production from the RT process as well as the polymerase extension step. Though not presented here, we have also determined that ingredients in UTM inhibit our RT-PCR reactions much more than VTM. This can explain why samples 1, 8, and 9 (low viral load samples in UTM) were not tested positive. These results suggest that the limit of detection under these conditions is around the Ct values of low 30s.

**Figure 4.**
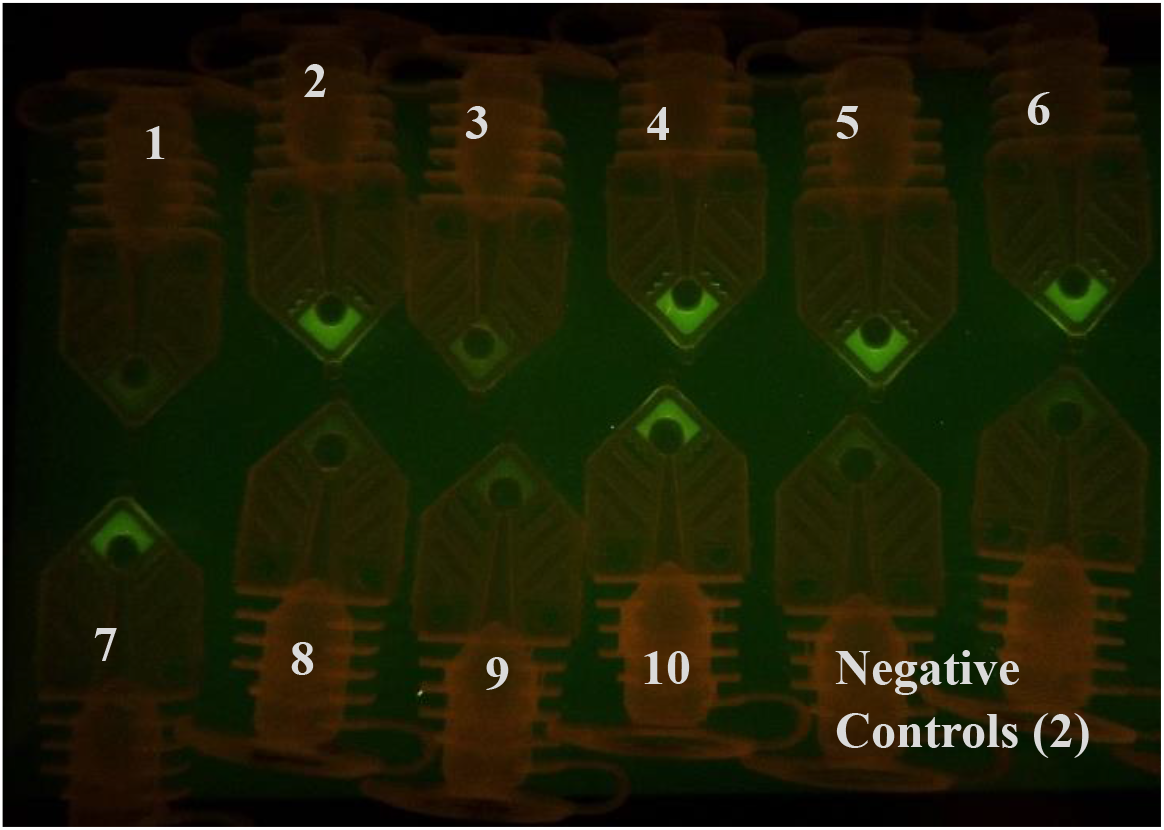
PCR tubes after a RT-PCR run using 3 μL of untreated transport media samples. Seven out of ten samples can be identified as positive COVID-19 patient samples by comparing results with negative controls. The RT-PCR run was completed in 12 minutes and the RT-PCR protocol was (90s/30s/40x(6s/9s)).

### Rapid RT-PCR detection of SARS-CoV-2 in heat-inactivated clinical samples

To reduce inhibition and improve sensitivity, a 10-minute, 95°C heat inactivation step was added to treat the samples. Three microliters of these samples were then added into RT-PCR master mix and perform the rapid test (N1). The photo taken after 40 cycles shows a significant increase in the fluorescence signal in 9 out of 10 samples (**Figure 5**). The signal intensity for sample 8, the sample with the highest threshold cycle value (Ct =35.96) among the 10 samples, was not different from the VTM-only negative control. The improved sensitivity from using heat inactivated samples for testing COVID-19 is similar to our results with flu sample testing (data not shown) and from others reported recently.^9^

**Figure 5.**
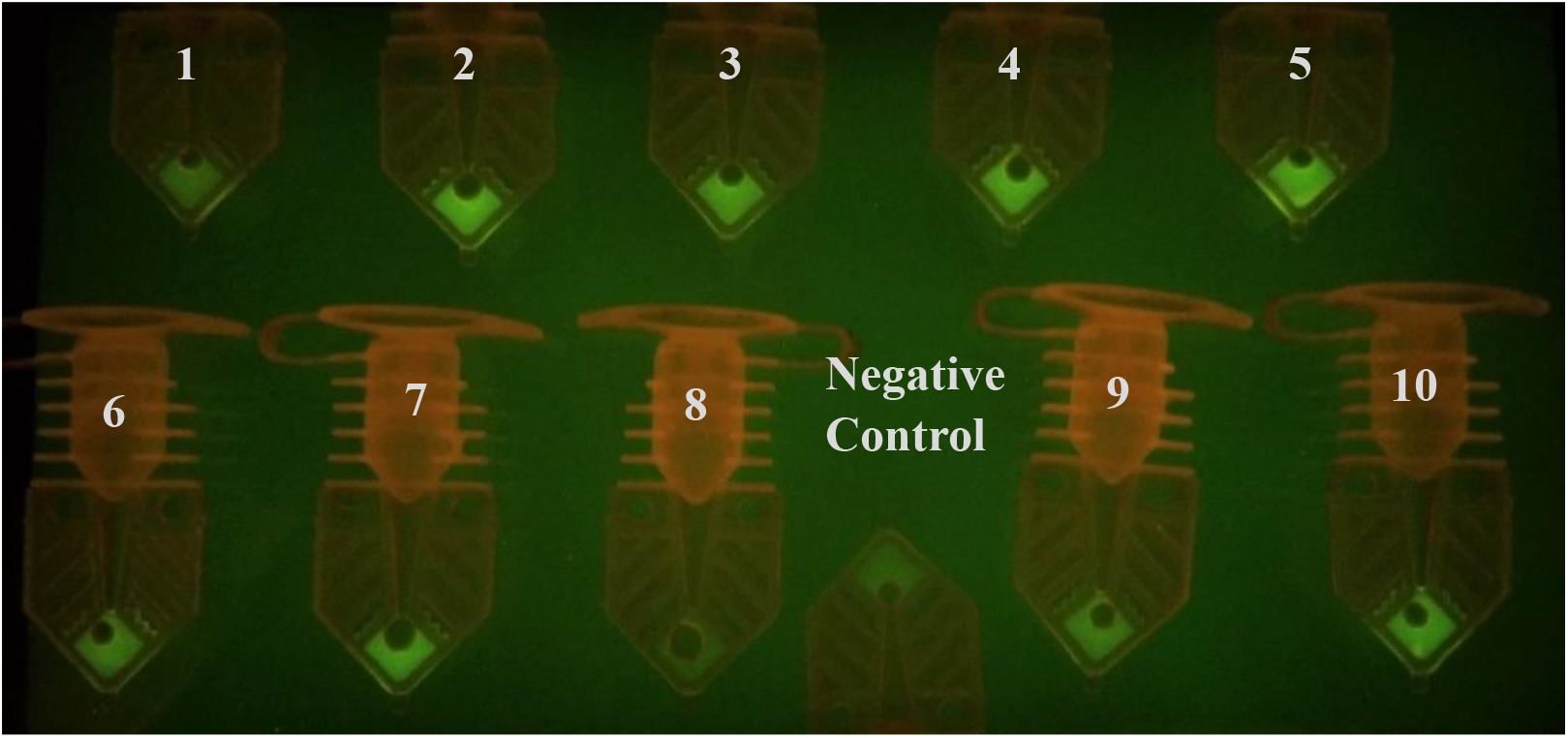
PCR tubes after a RT-PCR run using 3 μL of 10-minute, 95°C heat-inactivated samples. Nine out of ten samples can be identified as positive based on the fluorescence signal difference when compared to the negative control. The RT-PCR run was completed in 12 minutes (90s/30s/40x(6s/9s)).

### RT-PCR detection of SARS-CoV-2 with a 3-minute extraction protocol

We next tested whether we can use a rapid RNA extraction step to produce RNA templates for rapid RT-PCR from raw samples (no heat inactivation). We developed our own rapid magnetic particle-based extraction protocol using lysis buffer and two wash buffers before elution took place at 70 to 75°C. Three μL of the extracted templates was added to the RT-PCR master mix and tested in the water bath method. The photo taken after 40 cycles shows that we can call 10 out of 10 samples as N1 positive (**Figure 6**). The results suggest that a quick extraction step can improve sensitivity.

**Figure 6.**
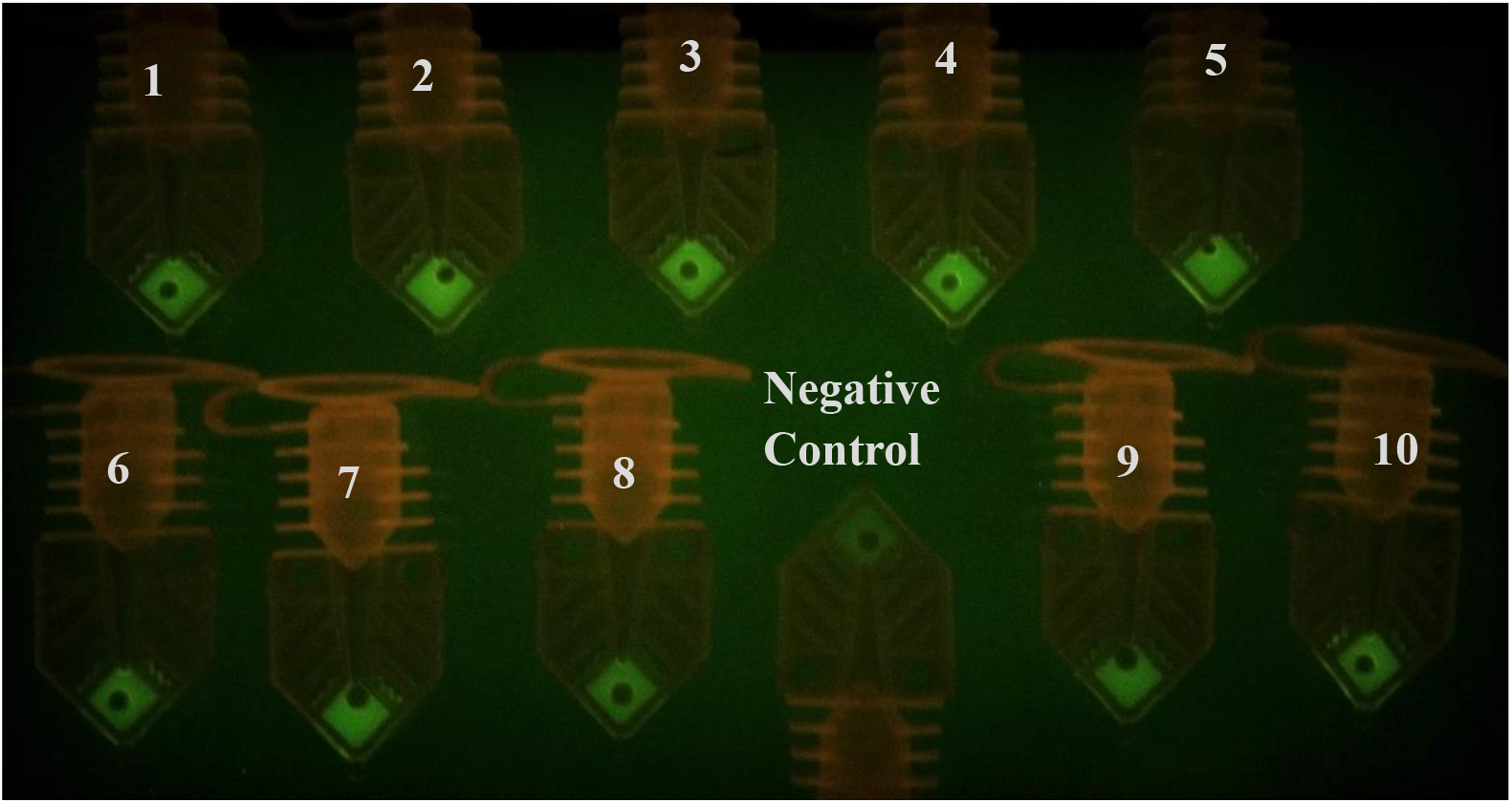
PCR Tubes after a RT-PCR run using 3 μL of extracted templates. Ten out of ten samples can be identified as positive based on the fluorescence signal difference when compared to the negative control. The RT-PCR run was completed in 12 minutes (90s/30s/40x(6s/9s)).

## Conclusions

Our post-RT-PCR photos of the PCR tubes show that water bath-based RT-PCR allows for 40 cycle RT-PCR reactions to finish in as little as 12 minutes. With a 3-minute RNA extraction protocol, we were able to get positive RT-PCR results with all ten COVID-19 positive clinical samples. Using raw samples after 10 minutes of heating, we were to detect 9 out of 10 samples using our approach. Using untreated samples with a 90-second reverse transcription step, we only were able to identify 7 out of 10 samples as positive. As expected, samples with a high viral load can easily be detected if the RT-PCR input uses raw or extracted templates. Samples with a low viral load or untreated COVID-19 positive samples are more likely to be missed if an extraction step is not used or when the PCR protocol is shortened when the presence of inhibitors affects the amplification efficiency.

For COVID-19 testing, using raw samples or minimal sample preparation steps may not significantly reduce the test sensitivity as most patients tend to have a higher viral load.^13^ Therefore, we report that the use of untreated samples can be a viable option during the COVID-19 pandemic. In addition, we can use a low-cost set up to allow fluorescence based end-point RT-PCR reactions to test for COVID-19 by running N1, N2, and RNase P RT-PCR reactions. This would allow locations with no access to real-time PCR thermal cyclers or even basic thermal cyclers to perform highly sensitive and specific gold-standard RT-PCR assays for COVID-19. The water baths are big enough to accept a 96-well PCR plate. The disadvantage of this approach is that a non-multiplexed reaction algorithm uses larger volumes of reagents. Also, if regular PCR tubes are used, the RT-PCR will take up to 30 minutes to complete.

## Materials And Methods

### Using a sous vide immersion heater to create a portable and low-cost alternative to circulating water baths used in laboratories

We have previously devised a low cost alternative to a thermal cycler using the thermos thermal cycler (TTC), a PCR method using thermos cans as insulated water baths to create a semi-automated cheaper alternative to conventional thermal cyclers to be used in low resource settings and small laboratories.^7,8^

For the current work, to achieve a steady circulating water system with consistent temperatures for denaturation and annealing/extension steps, we used two sous vide immersion heaters purchased at $99 each (Anova Culinary, 800 watt) to heat the water in two 6 quart clear food storage containers (Rubbermaid) to 55°C and 97°C, both of which are common PCR annealing and denaturing temperatures. The temperature of the 55 °C bath was lowered to 55°C during the reverse transcription step. The sous vide immersion heater is a medium-sized device that can be securely clamped onto the edge of the food storage containers. Its steady heating and water circulation functionality results in a well-maintained, even temperature throughout the bath for long periods of time. We were able to measure the temperature throughout the heating duration using a temperature data logger (HH147U, Omega Engineering, Norwalk, CT).

**Figure 7.**
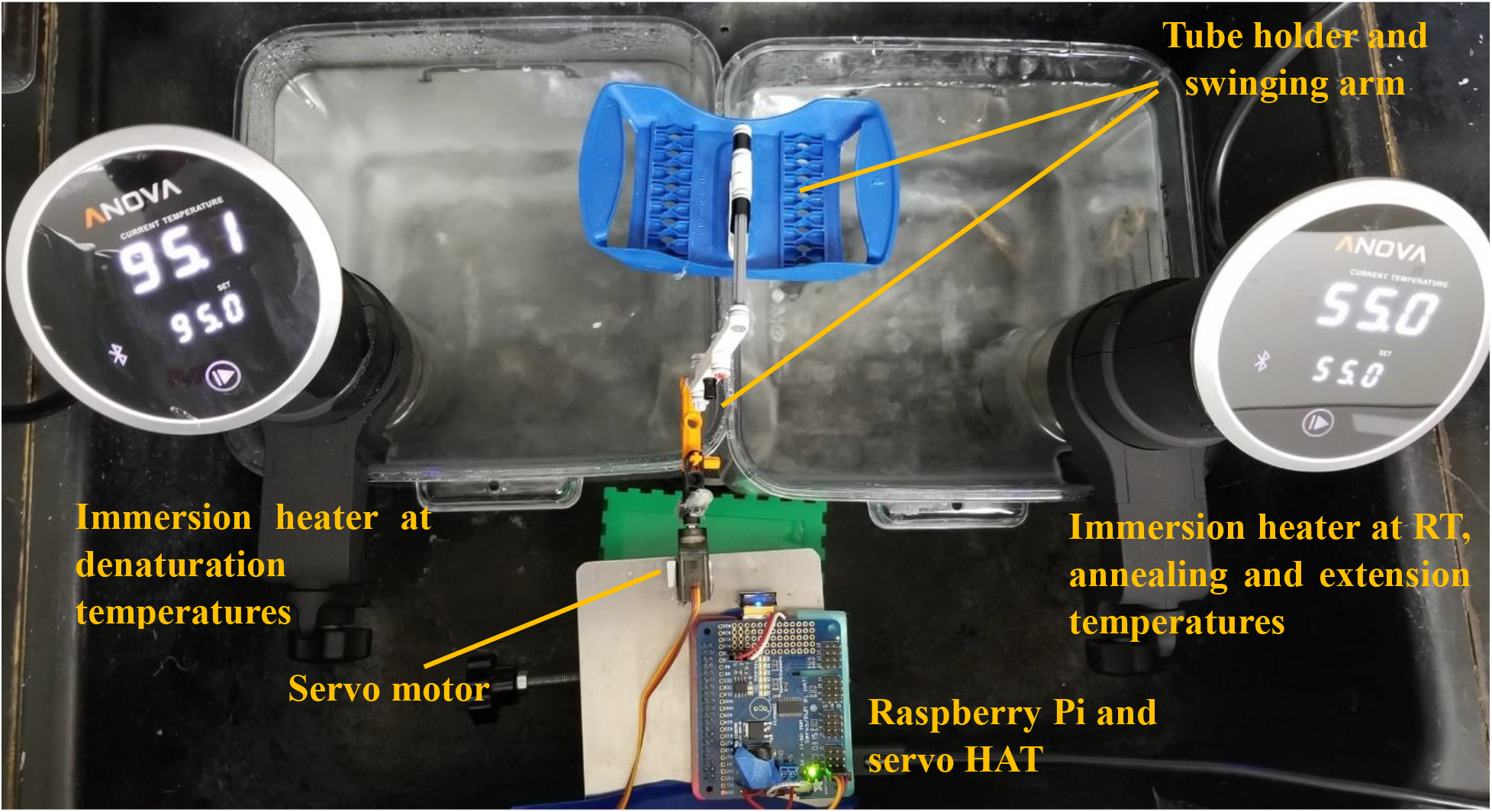
Water bath setup for SARS-CoV-2 detection using RT-PCR. Sous vide immersion heaters provided sufficient and consistent temperatures for RT, denaturation and annealing/extension steps. A Raspberry Pi controlled a servo motor that moved the PCR tubes between the baths with a cell phone app.

### Automation using a one-servo device

Automation of the thermal cycling reactions was achieved by programming a one-servo motor and devising a holder assembly with LEGO pieces to shuttle PCR tubes between the water baths (**Figure 7**). An arching movement was used to lift, transport, and lower the tubes between two baths in a single semi-circle motion. The tubes were contained in a hinged holder that would allow the tubes to remain upright throughout the movement phase. The mechanical portion was driven by a micro servo (MG90S Micro Servo Motor, Amazon) and was controlled by a microcontroller (Raspberry Pi Model A+) using a 16-Channel PWM / Servo HAT (Adafruit). Precise movement by the servo moved the arm back and forth between the two baths. The semi-circle movement of the arm, therefore, can move the PCR tubes in and out of the water. The entire arm assembly was mounted on a small lab-jack. A USB connector powered the servos and controller.

### Materials and Reagents

The COVID-19 positive clinical samples were obtained from the Medical College of Wisconsin (MCW) and consisted of de-identified nasopharyngeal specimens collected for non-research purposes. Specimens were defined as positive by a laboratory defined assay consisting of CDC primers extracted on an EMAG (Biomerieux, France) and tested on the 7500Dx (Applied Biosystems Fosters City, CA). Specimens were frozen within 24 hours of collection. Primer and probe sets for the SARS-CoV-2 RT-qPCR assay (Cat. No. 10006606) were purchased from Integrated DNA Technologies (Coralville, IA). The TaqPath 1-step multiplex master mix (Cat. No. A28525) was purchased from Thermo Fisher Scientific (Waltham, MA). The viral transport media (Cat. No. R99) used as negative controls was purchased from Hardy Diagnostics (Santa Maria, CA).

### Promega Maxwell extraction

The Promega Maxwell extraction was performed using AS1520 cartridge (Promega Corporation, Madison, WI). 60 μL of the samples from MWC was added to the cartridge and eluted in 60 μL of the elution buffer provided in the kit.

### Rapid extraction

We have used a rapid sample preparation protocol to reduce the sample-to-test time. 60 μL of the samples from MCW were added to 400 μL of lysis buffer and 15 μL of magnetic bead solution (NucliSENS Magnetic Particle Extraction Kit, bioMerieux, Durham, NC) and lysed for 1 minute. The beads were collected and washed in wash buffers 1 and 2 for 30 seconds each and eluted in 60 μL of elution buffer at 75°C for 1 minute.

### RT-qPCR reaction setup

TaqPath 1-step multiplex master mix was used for the RT-qPCR reaction with specific primers and probes tagged with FAM. For the detection of SARS-CoV-2 target genes, N1 and N2, the final concentration of primers was 500 nM and probes were 125 nM as per the CDC protocol (**Table 2**).^14^ Each PCR reaction was 20 μL with 3 μL samples/extracted template added. The typical run time for the Bio-Rad cycler to complete 50 cycles was 82 minutes. (**Table 3**)

**Table 2:**
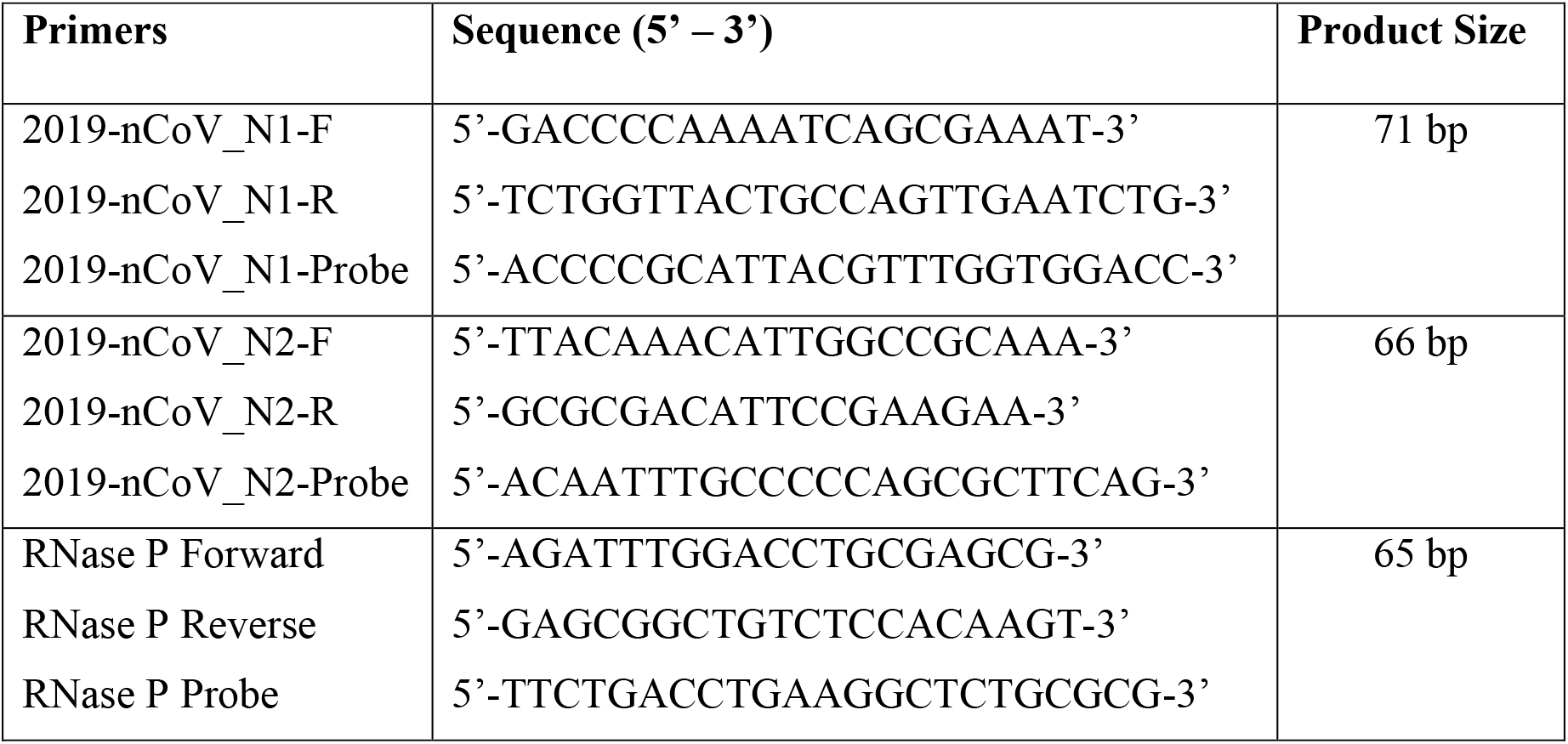
Primer and probe sequences

**Table 3:** Bio-Rad RT-qPCR setup

| Process | Temperature | Time | Cycles |
| --- | --- | --- | --- |
| UNG incubation | 25°C | 2 minutes | 1 |
| RT | 50°C | 15 minutes |  |
| Polymerase activation | 95°C | 2 minutes |  |
| PCR | 95°C | 3 seconds | 50 |
|  | 55°C | 30 seconds |  |

### Rapid PCR

Rapid PCR was conducted using two water baths maintained at 95°C and 55°C (**Table 4**). Reverse transcription was carried out for 90 seconds at 53°C and initial denaturation was carried out for 30 seconds followed by 40 cycles of PCR amplification with 6 seconds of denaturation at 95°C and 9 seconds of annealing/extension at 55°C. The fluorescence intensity of the reaction mix was analyzed after amplification. The total time for PCR was 12 minutes, including 2 minutes of RT and initial denaturation, and 10 minutes of PCR amplification for 40 cycles. It took 13.3 minutes for 45-cycle reactions (2 minutes of RT and initial denaturation and 11.3 minutes of PCR amplification). Protocols that deviated from these typical settings are noted in the results section. PCR tubes used in this study was Cepheid SmartCycler reaction tubes and thin-walled polypropylene PCR tubes (Cat. No. 16950, Sorenson Bioscience).

**Table 4:** Rapid RT-PCR setup

| Process | Temperature | Time | Cycles |
| --- | --- | --- | --- |
| RT | 53°C | 90 seconds | 1 |
| Polymerase activation | 95°C | 30 seconds |  |
| PCR | 95°C | 6 seconds | 40 or 45 |
|  | 55°C | 9 seconds |  |

### End-Point RT-PCR Fluorescence Analysis

After amplification by the water bath thermal cycler, PCR tubes were placed above a gel-viewing blue LED powered light box (Lonza Flashgel Dock or IO Rodeo Midi Blue LED Transilluminator) to confirm amplification through fluorescence intensity. Fluorescence across reactions was imaged using a smartphone camera (Note 8, Samsung Electronics) with an amber filter placed in front of the lens. We visually determined the RT-PCR result as positive or negative using the negative control as background fluorescence. Alternatively, software such as ImageJ can be utilized.

## ETHICAL STATEMENT

The intent of the work was for clinical method development as a response to the COVID-19 pandemic. In this work we used anonymized remnant material from samples that had been collected for clinical diagnostics of SARS-CoV-2. Our clinical specimens were provided by the Medical College of Wisconsin. These samples included oropharyngeal swabs in UTM and M6 VTM. These de-identified samples are not considered to be Human Subjects.

